# Functional analysis of PsbS transmembrane domains through base editing in *Physcomitrium patens*

**DOI:** 10.1101/2022.05.24.491476

**Authors:** Claudia Beraldo, Anouchka Guyon-Debast, Alessandro Alboresi, Fabien Nogué, Tomas Morosinotto

**Affiliations:** Dipartimento di Biologia, Università di Padova, Padova, Italy; Université Paris-Saclay, INRAE, AgroParisTech, Institut Jean-Pierre Bourgin (IJPB), 78000, Versailles, France

## Abstract

Plants exposed to light fluctuations are protected from photodamage by non-photochemical quenching (NPQ), a reversible mechanism that enables dissipation of excess absorbed energy as heat, which is essential for plant fitness and crop productivity. NPQ requires the activity of the membrane protein PsbS that, upon activation, interacts with antenna proteins, inducing their dissipative conformation. Here, we exploited base editing in the moss *Physcomitrium patens* to introduce *in vivo* specific amino acid changes and assess their impact on PsbS activity, targeting transmembrane regions to investigate their role in protein–protein interactions. This approach enabled the recognition of residues essential for protein stability and the identification of a hydrophobic cluster of amino acids with a seminal role in PsbS activity. This work provides new information on the PsbS molecular mechanism while also demonstrating the potential of base editing approaches for *in planta* gene function analysis.

## Introduction

Sunlight provides energy supporting the life of photosynthetic organisms through the activity of protein supercomplexes Photosystem (PS) I and II. In nature absorbed light often exceeds the metabolism capacity to use excitation energy, driving to the production of reactive oxygen species (Li et al., 2009). A feedback mechanism known as non-photochemical quenching (NPQ) drives the safe dissipation of excess excitation energy as heat, preventing photo-oxidative damage. NPQ was shown to be particularly important for plants growing in dynamic natural conditions with mutants showing reduced fitness in outdoor conditions (Kulheim et al., 2002). Heat dissipation of excitation energy, however, also represents an energy loss, reducing photosynthetic efficiency and negatively affecting productivity if activated constitutively (Dall’Osto et al., 2005). The optimized modulation of NPQ was indeed shown to improve biomass productivity in tobacco plants in field conditions (Kromdijk et al., 2016), demonstrating that the balance between photoprotection and efficiency is essential for optimal productivity (Alboresi et al., 2019) and can be improved in crops through genetics (Ort et al., 2015).

NPQ activation is triggered by the decrease of pH in the lumen of thylakoids, a condition typically occurring when light absorption is higher than cell metabolic capacity. NPQ activation requires a thylakoid membrane protein, PsbS (Li et al., 2000), that is protonated when lumen pH decreases (Li et al., 2002). PsbS is a member of the Light Harvesting Complex (LHC) superfamily, but in contrast to most of these proteins, it does not bind many chlorophylls and carotenoids (Fan et al., 2015). PsbS activity relies on the interaction with antenna proteins (LHC) (Betterle et al., 2009; Gerotto et al., 2015; Nicol et al., 2019) and consequently NPQ capacity is also largely affected in the absence of antennas (Lokstein et al., 1993; Nicol et al., 2019). PsbS does not appear to interact with only one specific antenna protein nor it strongly binds to a specific site PSII-LHC supercomplex (Caffarri et al., 2009; Kereïche et al., 2010; McKenzie et al., 2020).

*In vivo* interactions between PsbS and LHCs are essential for PsbS activity and these are only partially reproducible *in vitro* (Wilk et al., 2013; Nicol and Croce, 2021), impairing deep analyses of the protein structure and residues essential for activity. This makes *in vivo* studies essential to obtain deep knowledge on its structure and function, to open the possibility of identifying modified versions of the protein to modulate plant fitness and optimization of biomass production. Physcomitrella (*Physcomitrium patens*) was selected as a model organism for this study because PsbS in *P. patens* is active in NPQ (Alboresi et al., 2010), and in this species mutant generation is rapid and most tissues are haploid, making the phenotype assessment faster by avoiding the need of waiting for the progeny. Finally, methods for precise base editing have been set up and validated showing high editing efficiency (Guyon-Debast et al., 2021), making *P. patens* a suitable model for this kind of study. In this work, we employed advanced genome editing approaches based on CRISPR–Cas technology, cytosine base editor (CBE) and adenine base editor (ABE), to introduce specific nucleotide changes and targeted genetic diversity in the *PsbS* gene. Generation of various PsbS variants at the amino acid level enabled us to assess the impact of amino acids changes on PsbS *in vivo* activity, identifying key residues for protein structure as well as a cluster of hydrophobic amino acids influential on protein activity, demonstrating the power of the approach for precise genetic engineering and gene function analysis.

## Results and Discussion

In contrast to vascular plants, NPQ activation in the moss *P. patens* depends on two different proteins with additive contributions, PsbS and LHCSR(Alboresi et al., 2010; Gerotto et al., 2012). For this reason, *PsbS* mutagenesis was performed in *lhcsr1* KO plants to straightforwardly evaluate the impact of changes in the PsbS sequence and NPQ (Figure S1(Alboresi et al., 2010)). The mutagenesis specifically targeted the 4 transmembrane helices of PsbS, following the hypothesis that they play an important role as the sites of interaction with other integral membrane proteins, such as LHCs, which are essential for NPQ activity(Lokstein et al., 1993; Nicol et al., 2019). These transmembrane helices were targeted by designing 7 sgRNAs (Figure 1A, Table S1). *lhcsr1* KO plants were transiently transformed using a vector expressing one Cas9 nickase fused either to a cytosine or an adenine deaminase base-editor (CBE and ABE, respectively) together with one or multiple sgRNAs (Figure S2).

**Figure 1.**
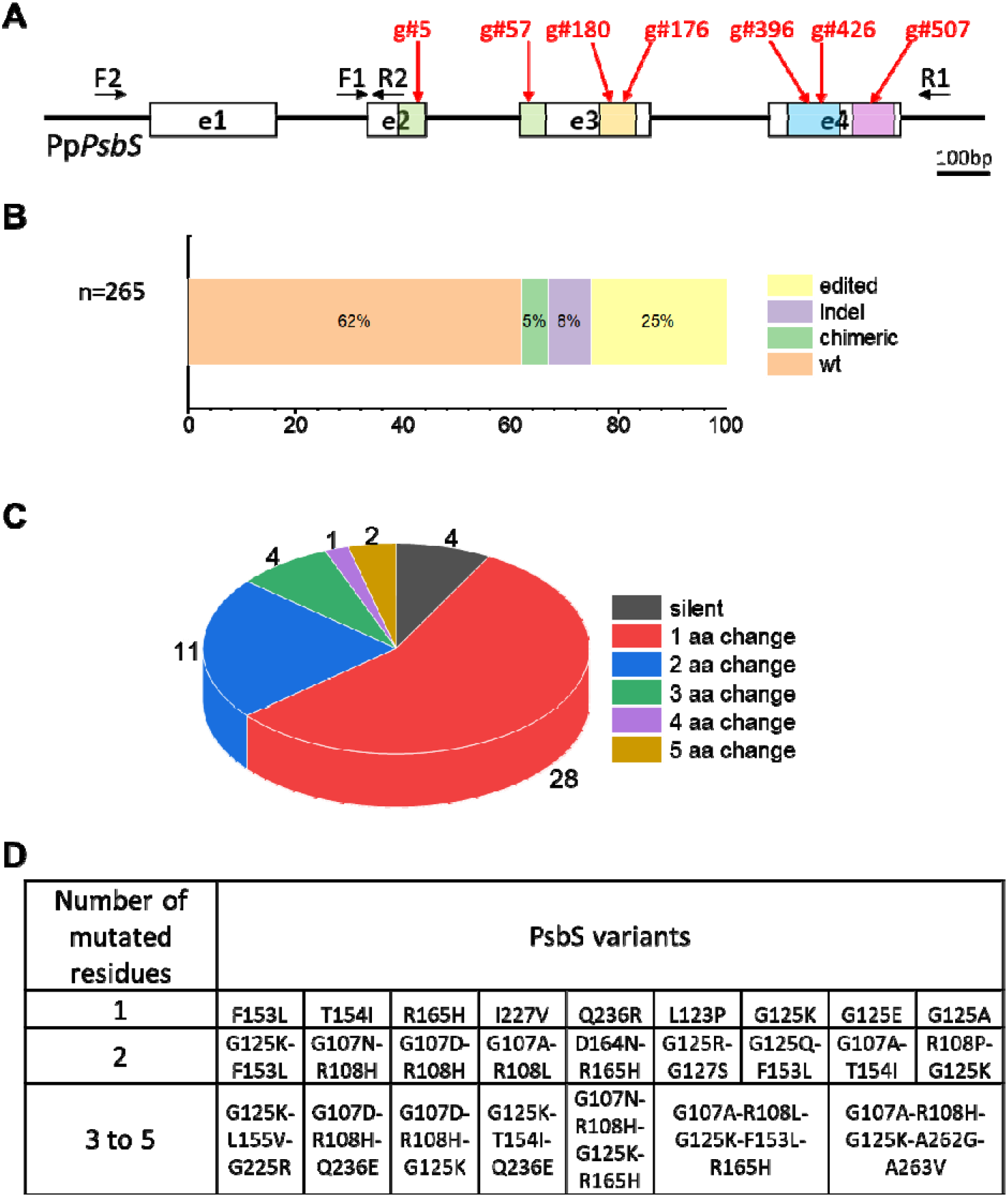
Multiplex base editing strategy for the generation of variants of the PSBS protein. (a) Structure of the *PpPsbS* gene and sgRNA positions. Boxes in white represent the exons, and black lines represent the introns. The coding sequences of the four transmembrane domains are indicated with colored boxes (green for TM1, yellow for TM2, blue for TM3 and purple for TM4). The seven sgRNA positions are indicated in red, and the primers used for PCR and sequencing are shown in black. (b) Results of *PpPsbS* gene sequencing of 265 transfected plants (multiplex and simplex strategy) using F1 and R1 primers. (c) Number of amino acid changes in the edited plants. (d) List of all PSBS-edited lines generated and phenotypically characterized.

Greater than 300 plants transfected by the CBE and ABE systems using multiplex or simplex strategies were phenotypically screened using a chlorophyll *a* fluorescence imaging apparatus to assess NPQ (Figures S3). The data showed that different lines had variable NPQ activity, demonstrating the power of the approach in generating functional variability (Figure S3C). Plants were subsequently amplified, starting from a small portion of tissue to ensure genetic homogeneity and avoid propagating possible chimeras.

All generated plants were genotyped by PCR, and the amplified products were sequenced. The *PsbS* gene of 265 plants (78% of generated plants) showed the expected size with no obvious chimerism or large deletions, which were otherwise observed in the other cases. Deletions were mostly detected in lines generated with a multiplex strategy, where simultaneous targeting of multiple sgRNAs can drive large deletions (Figure S4). PCR-amplified fragments with expected sizes were sequenced. Out of the 265 independent plants, 163 plants showed the *PsbS* WT sequence (62%), whereas 102 showed sequence modifications (38%). In total, 68 plants showed different combinations of edited bases (25%, listed in Table S2), whereas some short indels and chimerism were detected in 8% and 5% of plants, respectively (Figure 1B). Overall, the BE system was active in greater than one-third of plants, and 25 different variants of the PsbS protein with 1- to 5-amino acid changes were obtained, confirming that the approach was highly effective in generating a large set of genetic variants (Fig. 1C and Table S2). Although both multiplex and simplex strategies were effective in generating mutations in the expected regions, the methods yielded different results. Lines obtained using multiplex transformation exhibited greater variability than simplex-derived plants and had only 1-2 edited bases but with a much larger fraction of lines with WT sequences (Figure 1 and Table S2). The binding efficiency of each sgRNA, however, likely plays a major influence on the results. Thus, the simplex strategy was only performed for sgRNAs shown to be less effective in earlier multiplex experiments.

Functional analysis of all plants with the *PsbS* WT sequence showed NPQ equivalent to the parental line (Figure S5A), demonstrating an optimal consistency between genotype and phenotype. On the other hand, all plants with deletions in the *PsbS* gene showed impaired NPQ, which is similar to that noted for the *psbs* KO (Figure S5B). These results demonstrate that all of these edited bases caused complete protein inactivation, even the smallest ones missing only 9 amino acids (Figure S5C).

For all plants carrying one or more edited bases, the *PsbS* locus was sequenced again after a second round of subclone isolation (Figure 1D and Table S2) to confirm their genotype. In a few cases, base editing introduced a stop codon, and all these plants showed no detectable protein accumulation or NPQ activity (Figure S6). The latest stop codon found in the coding sequence was L237*, which causes the loss of the 4^th^ transmembrane helix (Figure S6). These results demonstrate that this region is essential for protein stability.

Taken together, the genetic variants showed modification of 19 amino acids over target protein regions (Figure 1). All targeted amino acids but two (L123 and T128) are fully conserved in the *PsbS* sequence of *P. patens* and those of other angiosperms, making the phenotypes observed translatable to other species (Figure S7, Table S3). All 9 PsbS variants isolated carrying 3 or more mutations showed complete loss of NPQ activity (Figure S8). This was always associated with a lack of accumulation of PsbS protein that was undetectable in all plants. This finding suggests that multiple mutations increase the probability of driving protein destabilization and, consequently, loss of activity (Figure S8).

Concerning all the other PsbS variants, in some cases (e.g., G125K, G125A, Figure S9), no significant alteration in protein activity or accumulation was noted, and the base-edited plant NPQ phenotype was indistinguishable from the WT. These results demonstrate that these residues have a minimal influence on protein activity. In other cases, mutations caused strong protein destabilization with a reduction in PsbS accumulation and consequent loss of NPQ activity. In L123P, the mutation into proline is expected to cause destabilization of the alpha-helix structure, destabilizing the overall protein folding (Figure S10). G125R-G127S plants showed lower PsbS accumulation and no measurable NPQ activity (Figure S10). Given that other single G125 mutants have no phenotypic alterations, this observation is attributable to G127, where the substitution with S can generate an electrostatic repulsion with E121 from TM1, destabilizing the protein (Figure S10). T154I also showed a decrease in NPQ activity correlated with a lower PsbS accumulation (Figure S11). T154 forms a hydrogen bond with F150 (Figure S11), which is involved in the formation of the hydrophobic cluster described below.

Mutations at D164 and R165 also caused destabilization of the protein with lower PsbS accumulation and a corresponding decrease in NPQ (Figure S12). An inspection of the PsbS structure showed that both residues are located at the end of TM2. The R165 carbon backbone interacts with F98 and K100, but these interactions are not expected to be altered by the mutations. The R165H mutation should instead alter polar interactions with G97 (Figure S12). D164 stabilizes polar interactions with K100 and R213 (Figure S12) in TM1 and 3, respectively. All residues are conserved in vascular plants (corresponding to residue numbering shown in Table S3), and these interactions are expected to be extremely relevant for the stabilization of protein structure, as confirmed by the strong impact of a mild modification such as D164N (Figure S12). Overall, these results point to electrostatic interactions in the stromal part of the protein as seminal for PsbS structural organization.

Mutations at the level of residue R108 from TM1 with various amino acid substitutions (R108P, R108H, R108L) also showed a strong impact on protein stability and activity (Figure S13). This arginine is part of a conserved transmembrane sequence typical of all LHC superfamily members, where it is involved in a salt bridge with the glutamic acid of another transmembrane helix important to stabilize the structure (Liu et al., 2004). Thus, the corresponding PsbS mutant in *A. thaliana* was already tested, showing consistent NPQ impairment (Schultes and Peterson, 2007). Structural investigation shows that R108 interacts with E208 on TM3 but also with A263 and N265 in TM4 (Figure S13). This is different from other LHCs, where transmembrane glutamic acid coordinates the Mg^2+^ of a chlorophyll. Given that the pigment is missing in PsbS, the extra partners for electrostatic interactions are derived from TM4 residues. This finding also points to different structural protein organizations in PsbS and LHCII that could be at the base of their different functions.

Q236R caused a lower accumulation of PsbS than in WT plants and a corresponding decrease in activity (Figure S14). Q236 is located in H2 and forms polar interactions with K231 and S222 in TM3 (Figure S14) that are likely altered by Q236R, causing protein destabilization and highlighting the relevance of this polar interaction for protein stability. All residues’ polar interactions identified as essential for PsbS protein stability are summarized in Figure 2.

**Figure 2.**
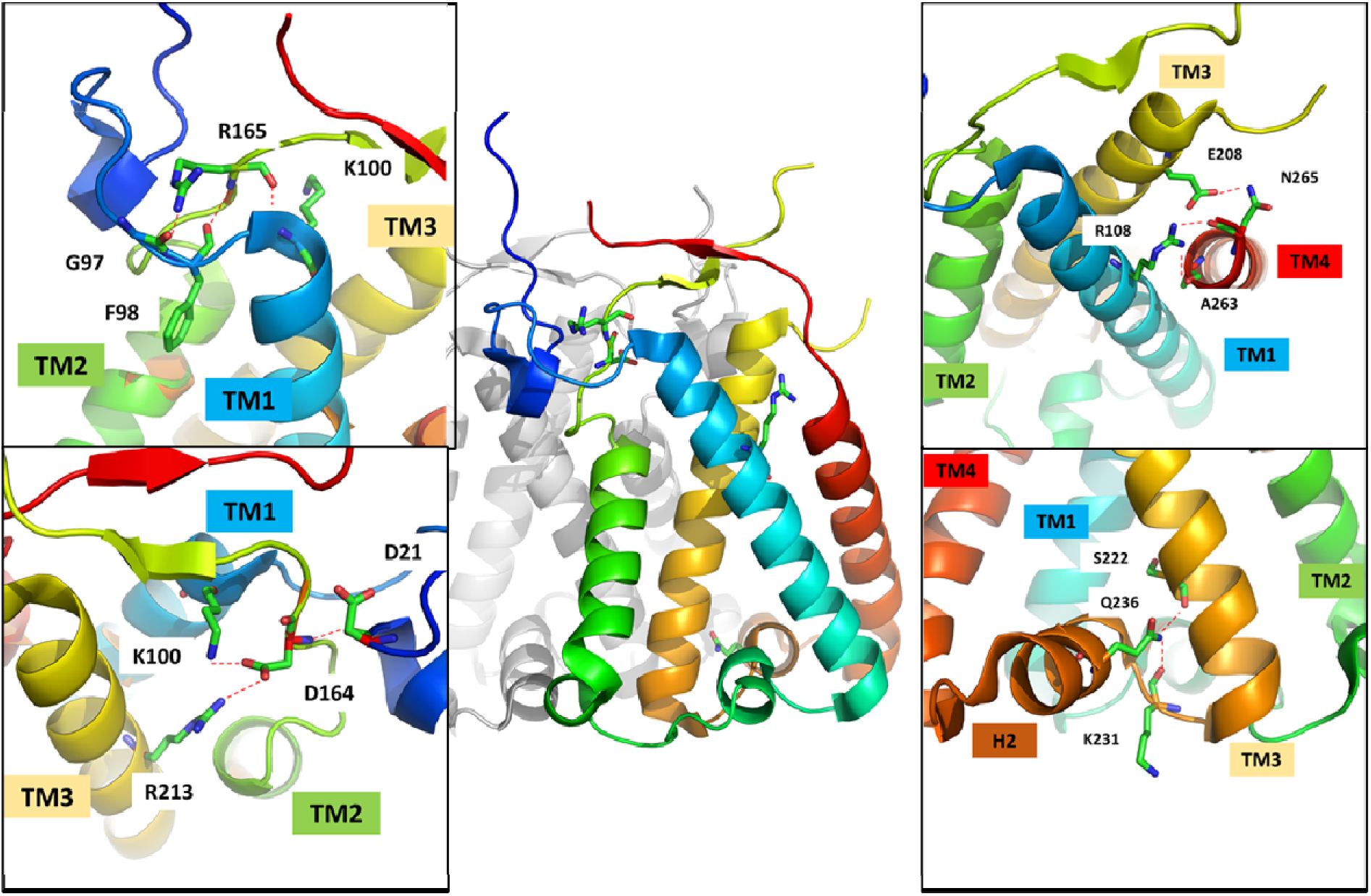
Overview of polar interactions stabilizing the PSBS structure identified by mutational analysis. Residues are shown as sticks, polar interactions with the dashed line. Nitrogen and oxygen atoms are colored blue and red, respectively. Different transmembrane helices (TM) are shown in different colors.

Another set of mutants showed altered NPQ activity, whereas PsbS accumulation was maintained, suggesting an alteration in protein activity. The F153L mutation caused an altered NPQ, as confirmed by other mutants carrying the same mutation in combination with changes in G125, which was already shown to have no impact on activity (Figure 3). Given that both leucine and phenylalanine have hydrophobic properties, the reduction in the activity of the F153L mutant is attributable to the absence of the aromatic ring, which may be involved in hydrophobic interactions. F153 is in the second transmembrane helix. In the PsbS structure, F153 interacts with n-nonyl-β-D-glucopyranoside (BNG), the detergent used for protein isolation before crystallization. F153 (TM2) is also close to F149, F150 (TM2), V106 (TM1), F98 (TM1) and L162 (TM2), and all hydrophobic residues (Figure 3) are well conserved in different organisms, suggesting a possible functional role (Figure S7) beyond the hydrophobicity required for membrane localization. This cluster of amino acids protrudes from the protein, points toward the membrane (Figure 3) and is not involved in intermolecular interactions within the PsbS dimer. This finding suggests that *in vivo*, these residues could interact with other hydrophobic components present in the membrane, such as lipids or antenna proteins, and that these interactions may indeed have an important functional role (Figure 3).

**Figure 3.**
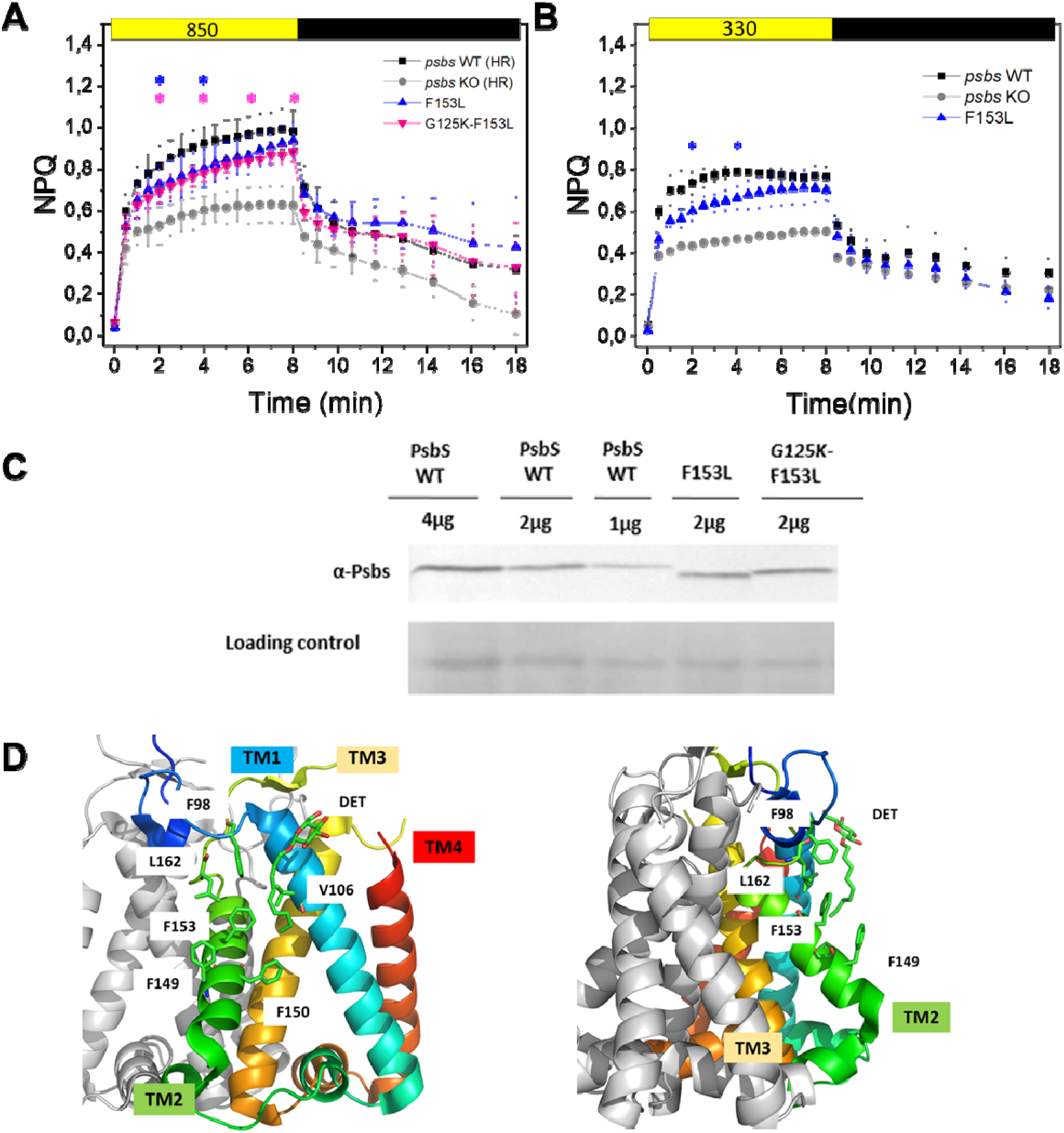
Analysis of the hydrophobic cluster. A-B) NPQ of F153L (blue triangles) and the G125K-F153L mutant (pink triangles) compared with the *psbs WT* line (black circles) and the *psbs* KO line obtained with homologous recombination (gray circles). Two different actinic lights were used: 850 mmol photons m^−2^ s^−1^ (A) and 330 mmol photons m^−2^ s^−1^ (B). Data represent average values ± SD, n=3. C) Immunoblot analysis evaluating PsbS accumulation in F153L mutants compared with the *psbs* WT line. D) Overview of the hydrophobic cluster. Residues are shown as sticks, polar interactions with the dashed line. Nitrogen and oxygen atoms are colored blue and red, respectively. Different transmembrane helices (TM) are shown in different colors.

In conclusion, the base editing approach was successful in generating a large set of plants with variable alterations in the PsbS sequence and carrying 1-5 amino acid variants. This enabled us to obtain valuable new information on PsbS, identifying several polar interactions essential for protein stability and pointing to differences in the structural organization in comparison to LHC. A cluster of hydrophobic amino acids with a potential role in protein–protein interactions was also identified, opening the way to the identification of elusive PsbS interactions essential for its biological activity. The work here was performed in *P. patens*, which provided advantages in the speed of generation, selection, and characterization of plants. On the other hand, all results are translatable to vascular plant species given the high conservation of the protein sequence.

## Materials and Methods

### Molecular cloning

Guide RNA (sgRNA) sequences specific to the *PsbS* gene (Pp3c20_23430) were chosen using the web tool CRISPOR 4.97 (Concordet & Haeussler, 2018). Expression cassettes sgRNA#5, sgRNA#57, sgRNA#176, sgRNA#180, sgRNA#396, sgRNA#426 and sgRNA#507, including the promoter of the *P. patens* U6 snRNA (Collonnier et al., 2017), the 5’-G-N(19)-3’ guide sequences targeting the *PsbS* gene and the tracrRNA scaffold, were synthesized by Twist Bioscience (San Francisco, California, USA; Table S1). Expression cassettes were subcloned into the pDONR207-NeoR vector(Guyon-Debast et al., 2021) by Gateway™ BP reaction (Invitrogen) as previously described to yield psgRNA#5, psgRNA#57, psgRNA#176, psgRNA#180, psgRNA#396, psgRNA#426 and psgRNA#507. For the CBE and ABE systems, we used the plasmids pnCas9-CBE1 (pAct-nCas9-CBE) and pnCas9-ABE1 (pAct-nCas9-ABE), as previously described (Guyon-Debast et al., 2021).

### Moss culture and transformation

Plants were grown on PpNH_4_ medium (PpNO_3_ medium supplemented with 2.7 mM NH_4_-tartrate) in growth chambers set at 60% humidity with 16 hours of light (quantum irradiance of 40 μmol m^−2^ s^−1^) at 24 °C and 8 hours of dark at 22 °C.

Moss protoplast isolation and transfection were performed as previously described(Charlot et al., 2022). Protoplasts were transfected with a total of 21 μg of circular DNA divided as follows: 7 μg of each base editor (pnCas9-CBE1 and pnCas9-ABE1 plasmids) and a mix of 7 μg of sgRNA plasmids for multiplex or 10 μg of the base editor (pnCas9-CBE1 or pnCas9-ABE1 plasmid) and 10 μg of the sgRNA plasmid (psgRNA#5, psgRNA#176, psgRNA#180, psgRNA#396 or psgRNA#426). After protoplast regeneration, plants were transferred to PpNH_4_ supplemented with 50 mg/L G418 (Duchefa) to select clones that were transiently transfected (Figure S2). A total of 189 transfected plants were selected using the multiplex strategy. To produce some additional specific genetic combinations, 151 transfected plants were selected using the simplex strategy.

### PCR and sequence analysis of the edited plants

For PCR analysis of transiently transfected plants, genomic DNA was extracted from 50 mg of fresh tissue as previously described(Lopez-Obando et al., 2016). Molecular analysis was based on PCR genotyping using primers surrounding the targeted locus, F1 and R1 primers. Sanger sequencing (Genoscreen, Lille, France) of the *PsbS* gene was performed on 265 plants showing no large deletion or obvious chimerism. The exon 1 sequence was also checked using the F2 and R2 primers for the 67 edited plants. The PCR primers used in this study are listed in Table S2.

### In vivo Chl fluorescence kinetics

For screening purposes, the *in vivo* chlorophyll fluorescence signal was monitored at room temperature with an imaging instrument, FluorCam 800-C (Photon System Instruments). Approximately one-month-old plants were exposed after a dark adaptation to actinic light (850 μmol photons m^−2^ s^−1^) for 4 minutes and then left to recover for an additional 4 minutes and 12 seconds in the dark.

After selection, the NPQ phenotype was assessed by the *in vivo* chlorophyll fluorescence signal at room temperature with a Dual-PAM-100 fluorometer (Walz, Germany) in protonemal tissues grown for 10 days in PpNO_3_ medium under control growth conditions (40 μmol m^−2^ s^−1^). Before measurements, plants were dark-acclimated for 40 minutes. For induction kinetics, actinic light was set to 850 (saturating actinic light) or 330 μmol photons m^−2^·s^−1^. PSII parameters were calculated as follows: Fv/Fm as (Fm - Fo)/Fm and NPQ as (Fm - Fm′/Fm′. Data are presented as the mean ± SD of at least 3 independent experiments.

### Total protein extracts, SDS page, and Immunoblotting

For Western blot analysis, 6 independently transfected plants (#11, 115, 154, 162, 177, 190) showing the *psbs* WT sequence were used as a reference given that PsbS WT from transfected plants had the same PsbS accumulation as the parental line. In addition, 7 independently transfected plants with deletions in the *psbs* sequence (# 18, 56, 81, 142, 143, 201, 202) were used as a reference given that lines with deletions in PsbS exhibited no PsbS accumulation, which is similar to that noted for the *psbs* KO line generated from homologous recombination (Figure S5). Total extracts were obtained by grinding tissues in sample buffer before SDS/PAGE. For immunoblotting analysis, after SDS/PAGE, proteins were transferred to nitrocellulose membranes (Pall Corporation) and detected with a specific homemade polyclonal PsbS antibody ((Gerotto et al., 2015)). The Chl a/b and Chl/Car ratios were obtained by fitting the spectrum of 80% acetone pigment extracts with spectra of the individually purified pigments(Croce et al., 2002).

## Funding

French National Research Agency (ANR11-BTBR-0001-GENIUS, FN).

Saclay Plant Sciences-SPS (ANR-17-EUR-0007, FN).

## Author contributions

Conceptualization: FN, AA, TM

Methodology: AA, FN, AGD

Investigation: CB, AGD, AA

Visualization: CB, AA, TM

Supervision: AA, FN, TM

Writing—original draft: TM

Writing—review & editing: CB, AGD, AA, FN

## Data and materials availability

All data are available in the main text or the supplementary materials.

