## Supplementary material for "Functional analysis of PsbS transmembrane domains through base editing in *Physcomitrium patens*": Supplemantal Figures and Tables

**This PDF file includes:**

Supplementary Text
Figs. S1 to S14
Tables S1 to S3

**Fig. S1.**

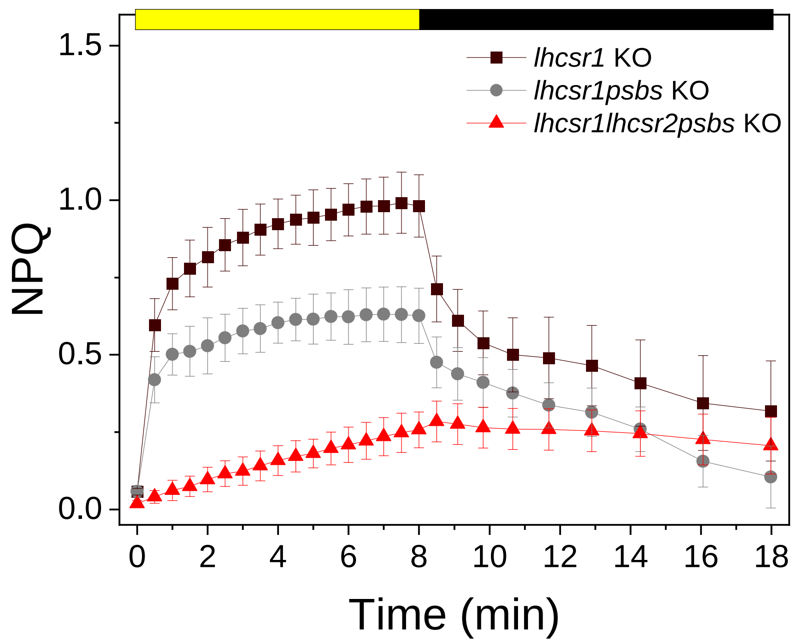


**Figure S1. NPQ induction in *P. patens* reference lines.** NPQ activation was determined for dark-adapted plants during 8 min of illumination at 850 µmol photons $m^{-2}$ $s^{-1}$ and a subsequent dark period of 10 min for the following genotypes, *lhcsr1 KO* (black squares), *lhcsr1psbs KO* (grey circles) and *lhcsr1lhcsr2psbs KO* (red triangles) *lhcsr1 KO* plants were selected as parental line to assess PsbS activity. Values represent the mean ± S.D. (n = 4 biological replicates).

**Fig. S2.**

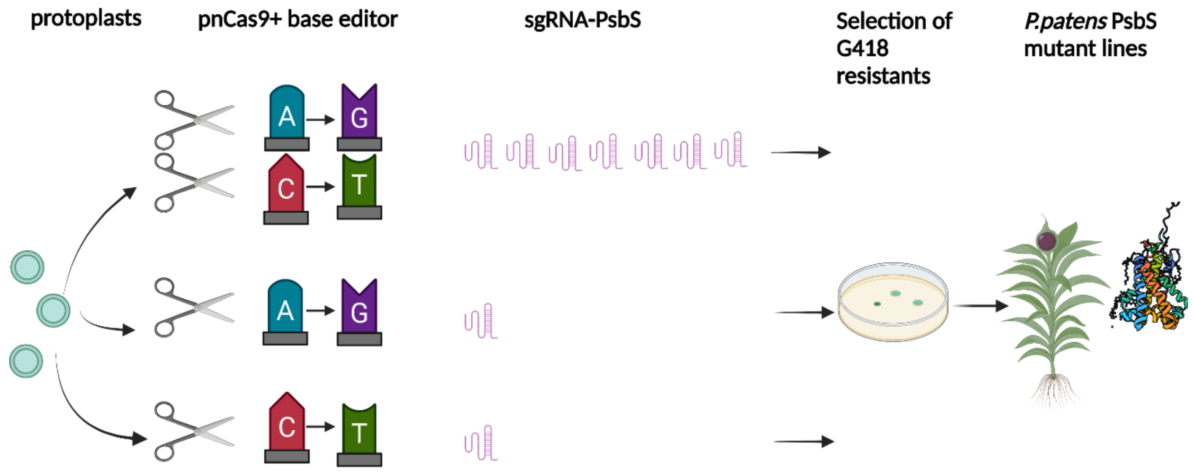


**Figure S2. Scheme of base editing approach using multiplex and simplex strategy.** A) Protoplasts of *lhcsr1* KO plants were transfected with pnCas9 (represented as scissors) + ABE and/or CBE which causes A-G or C-T conversion, respectively, and 1 or 7 plasmids expressing sgRNA targeting *PpPsbS* gene. The sgRNA were subcloned into vectors also containing the resistance to G418. After transfection, G418 resistant clones were transiently selected and different PsbS edited variants were finally propagated. B) The table reports the list of the three different combinations of plasmids used to transfect *lhcsr1* KO protoplast tested. Two different base editors, cystosine base editor and adenine base editor fused with nCas9 (cas9 nickase) were used with a combination of 7 sgRNAPsbS (multiplex transfection) or with a single sgRNA-PsbS (simplex transfection).

**Fig. S3.**

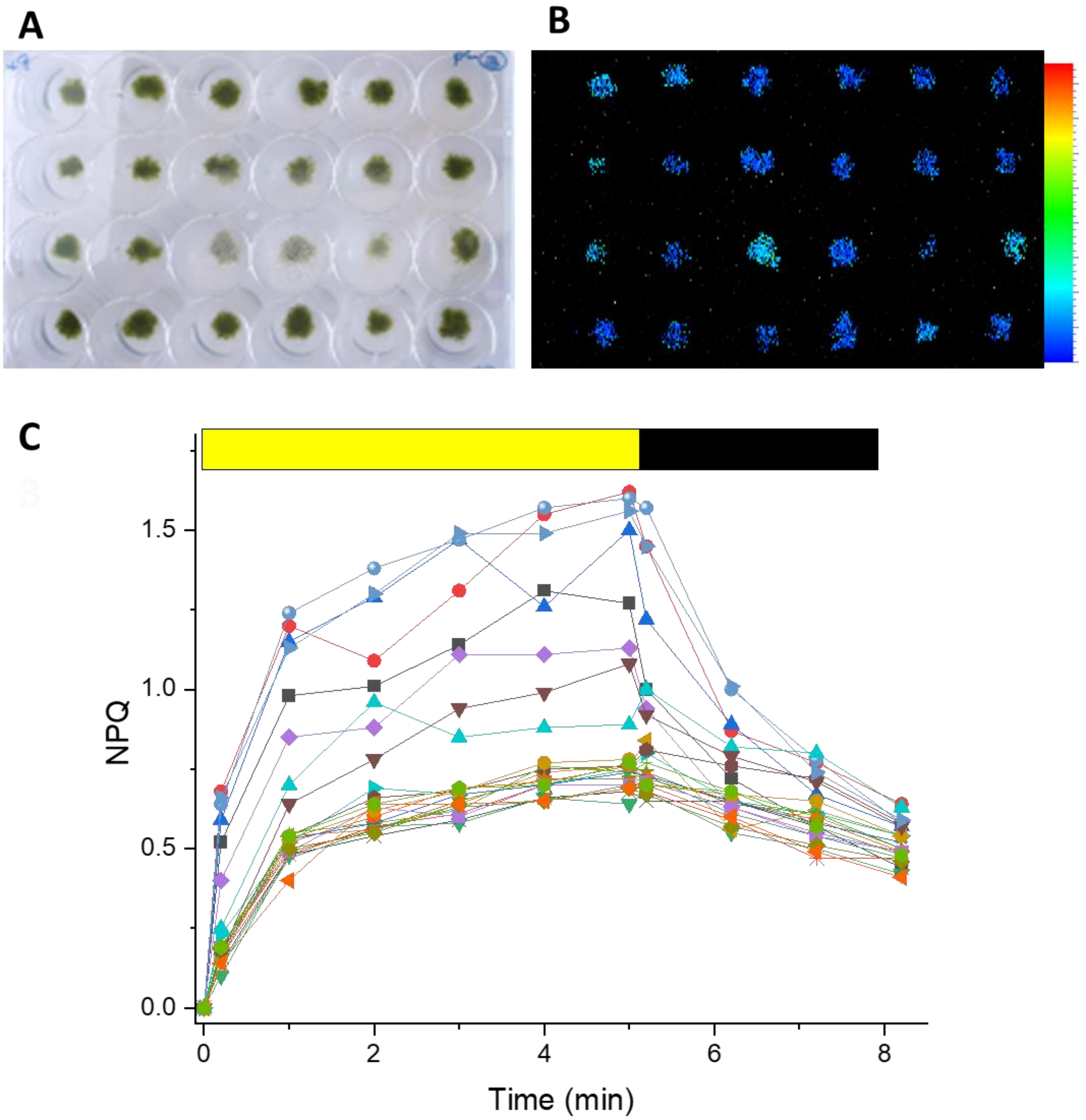


**Figures S3. Screening of NPQ capacity in edited plants.** Multiple independent lines obtained from transfection using multiplex and simplex strategies (A) were measured with a fluorescence imaging apparatus to quantify NPQ. In B) NPQ levels are represented as false colours,. In C) example of data obtained with 5 minutes of actinic light followed by 3 of dark relaxation. Independent lines are shown in different colours and symbols.

Type or paste caption here. Create a page break and paste in the Figure above the caption.

**Fig. S4.**

**A**

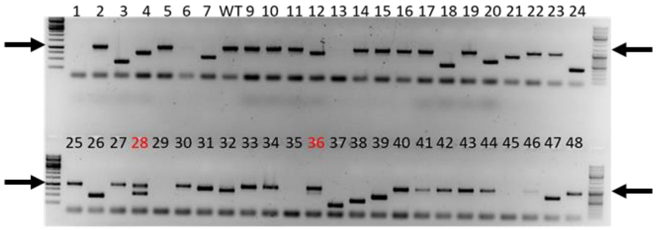


**B**

| Nature of plants | Multiplex | Simplex | Total |
|---|---|---|---|
| Expected PCR Size | 114 | 151 | 265 |
| Deletion | 62 | 0 | 62 |
| Chimeric | 13 | 0 | 13 |
| Total plants with positive PCR | 189 | 151 | 340 |

**C**

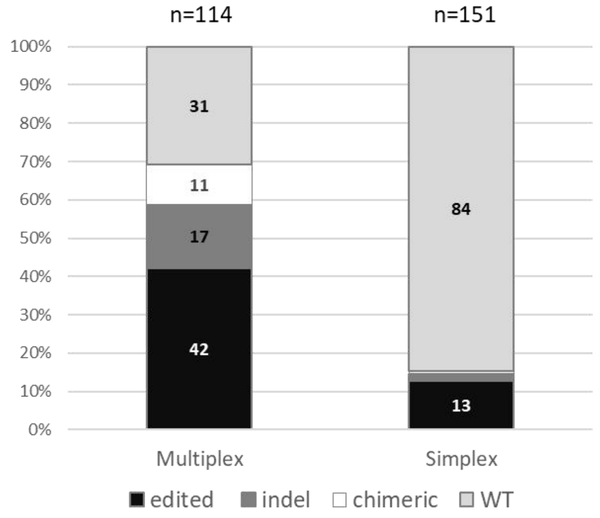


**Figure S4. Molecular analysis of transfected plants.** (a) Example of PCR genotyping agarose gel for Multiplex strategy using F1/R1 primers. Expected size amplicons are indicated with black arrows, chimeric plants are indicated in red. These by-products were only detected using multiplex system, large deletions are due to simultaneous targeting of multiple sgRNAs. (b) Results of PCR genotyping for multiplex and simplex strategy. (c) Results of *PpPsbS* gene sequencing for both strategies.

**Fig. S5.**

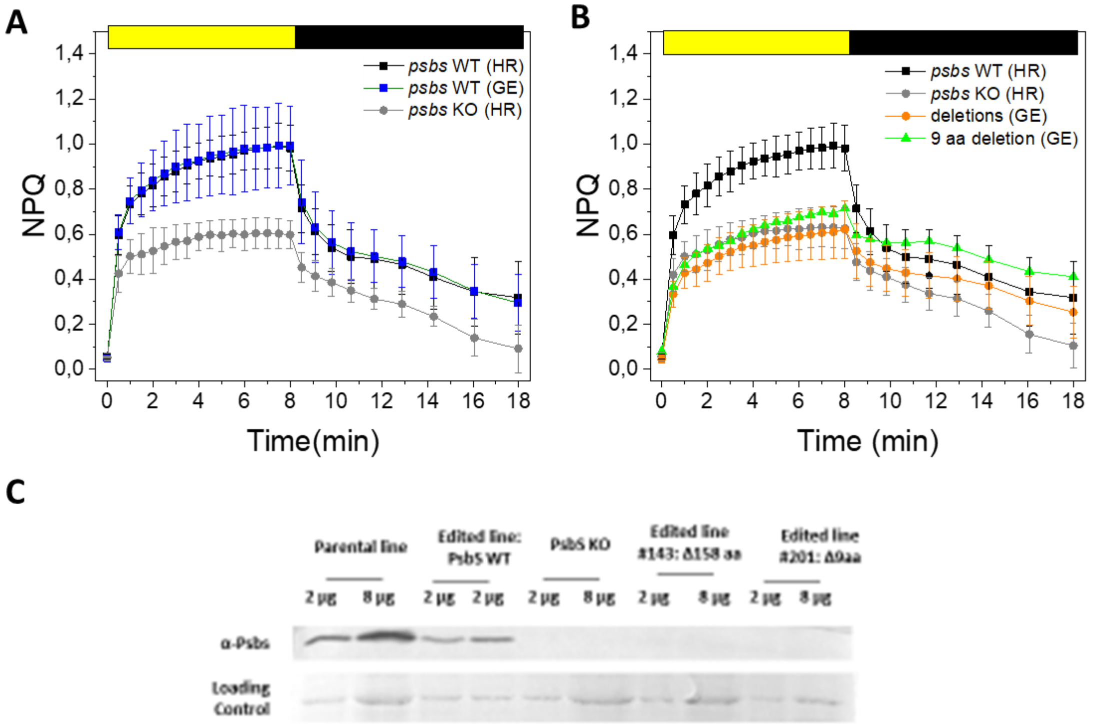


**Figure S5. NPQ of lines with WT PSBS and deletions.** A) NPQ averaging 6 independent lines obtained by base editing and showing WT PsbS sequence (blue squares) is compared with parental line (black squares) and *psbs* KO obtained from homologous recombination[1] (grey circles). B) Average NPQ of 7 independent lines showing deletions in PsbS (orange circles) compared with parental line (black squares) and *psbs* KO obtained from homologous recombination (grey circles). NPQ from Line #201, the one with the smallest deletion is also reported (green triangles). All measurements were performed for at least 3 biological replicates. C) WB analysis of different lines originating from editing experiments (GE) having WT sequence (#177 and #115) or deletions (#81 and #201) are compared with parental line and *psbs* KO obtained from homologous recombination. All lines have a *lhcsr1* KO background.

**Fig. S6.**

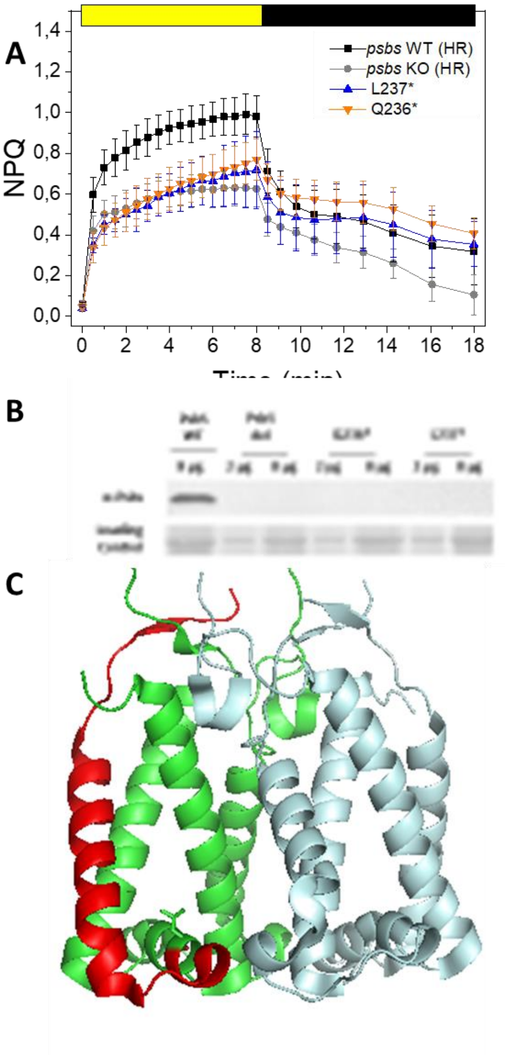


**Figure S6. Role of C terminal on PSBS structure.** A) NPQ phenotype of L237* (line #80- blue triangles) and Q236* (lines #63, #163- orange triangles) compared to the parental line (black squares) and *psbs* KO (grey circles). For all plants measurements show average of at least 3 independent biological replicates. B) WB against PsbS comparing *psbs* WT line resulting from genome editing, line with deleted PsbS and two lines with the latest stop codon (Q236* and L237*). C) Structure of PsbS showing in red the fraction of C terminal that was lost in L237*. Two Monomers are shown in green / red and grey.

**Fig. S7.**

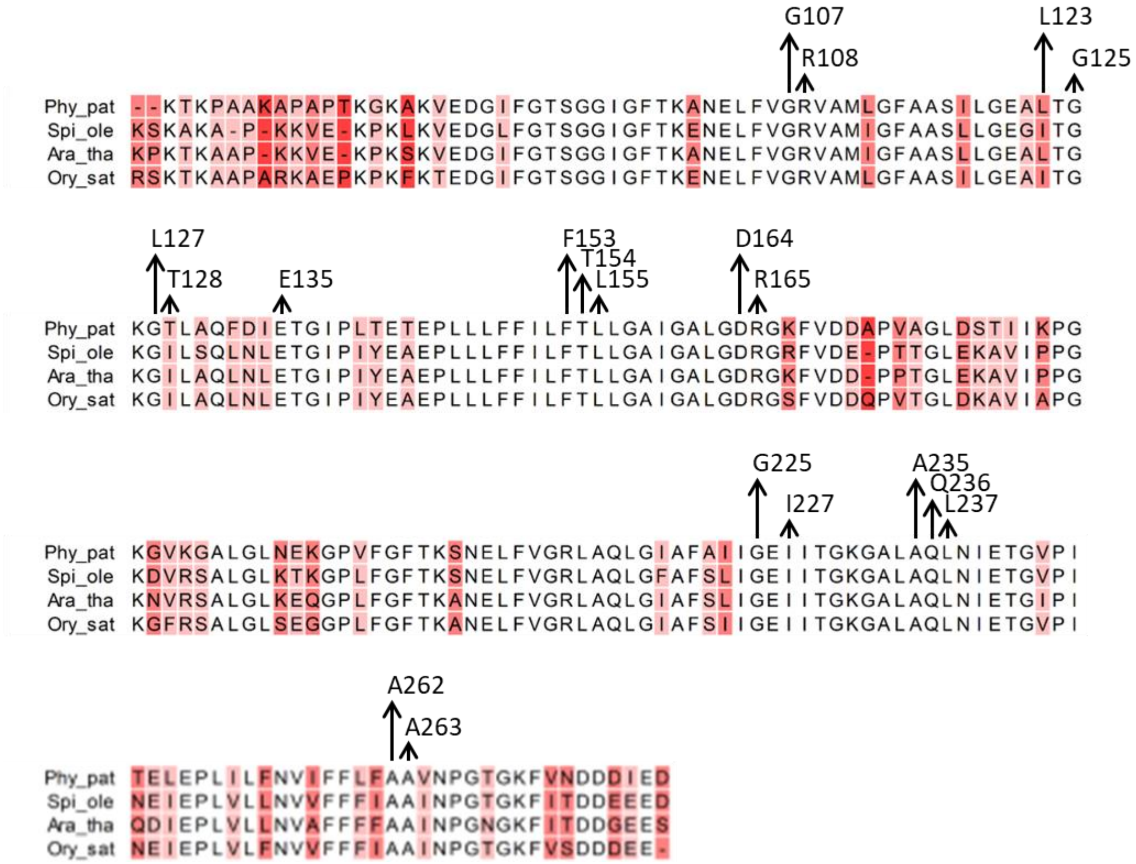


**Figure S7. Sequence alignment of PsbS proteins.** *Physcomitrium patens* protein (Uniprot code A9TJ15) is labeled as Phy_pat, *Spinacia oleracea* protein (Uniprot code Q02060) is labeled as Spi_ole, *Arabidopsis thaliana* protein (Uniprot code Q9XF91) is labeled as Ara_tha and *Oryza sativa* protein (Uniprot code Q943K1) is labeled as Ory_sat. Residues edited in *P. patens* PsbS protein are highlighted with an arrow and labeld with the number corresponding to aminoacid position. White background corresponds to full conservation of the residues among the four sequences, colored background highlights residues that are not conserved.

**Fig. S8.**

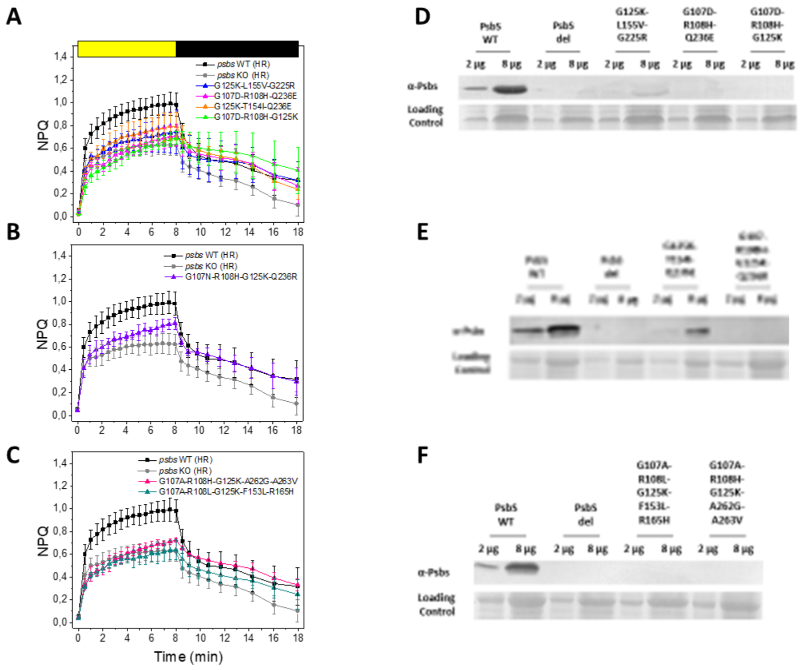


**Figure S8. Impact of multiple aminoacid changes on NPQ and PSBS accumulation**. NPQ kinetics of mutants lines with 3 (A), 4 (B) or 5 (C) aminoacids changes in PsbS sequence. In all graphs *psbs* WT (black squares) and *psbs* KO (grey circles) are also shown as reference. Mutants are represented as triangles of different colours, as detailed in the legend. For all plants measurements show average of at least 3 independent biological replicates. Western blotting monitoring PsbS accumulation in lines with 3 (D-E), 4(E) and 5 (F) aminoacids changes in comparison with *psbs* WT line and line with deletions in PsbS sequence.

**Fig. S9.**

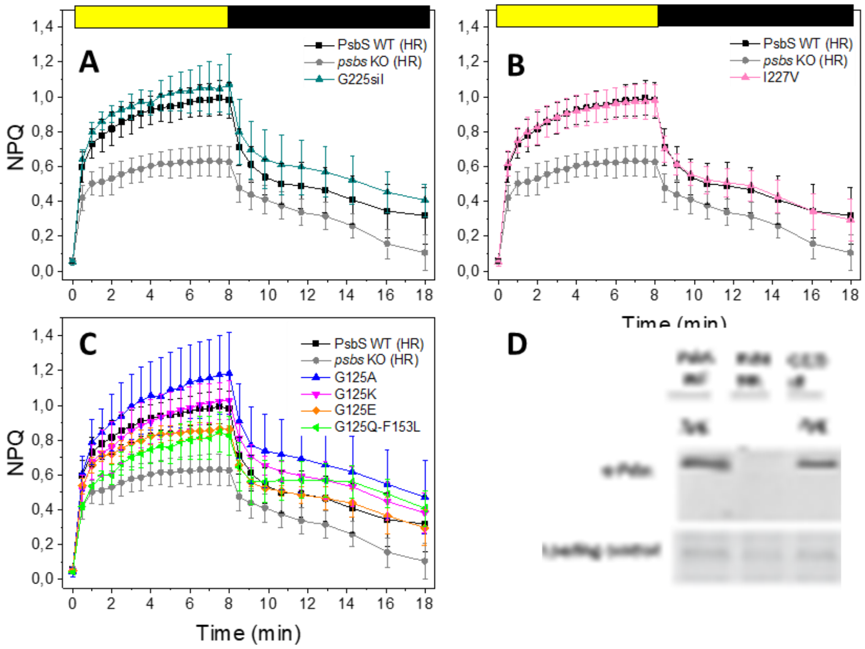


**Figure S9. Mutants with no effect on NPQ activity.** NPQ kinetics of different mutants lines (A) G225sil (line#102). (B) I227V (lines#195, #203, # 204), (C) G125A (line#129) /K (lines#23, #85, #147, #)/E (lines#14, line#85). The double mutant G125Q-F153L is also added (line#133). In all graphs *psbs* WT (black squares) and *psbs* KO (grey circles) are shown as reference. Mutants are represented as triangles of different colours, as detailed in the legend. For all plants measurements show average of at least 3 independent biological replicates. D) Example of Western Blotting analysis comparing G225sil PsbS variant with *psbs* WT line and line with deletions in PsbS sequence.

**Fig. S10.**

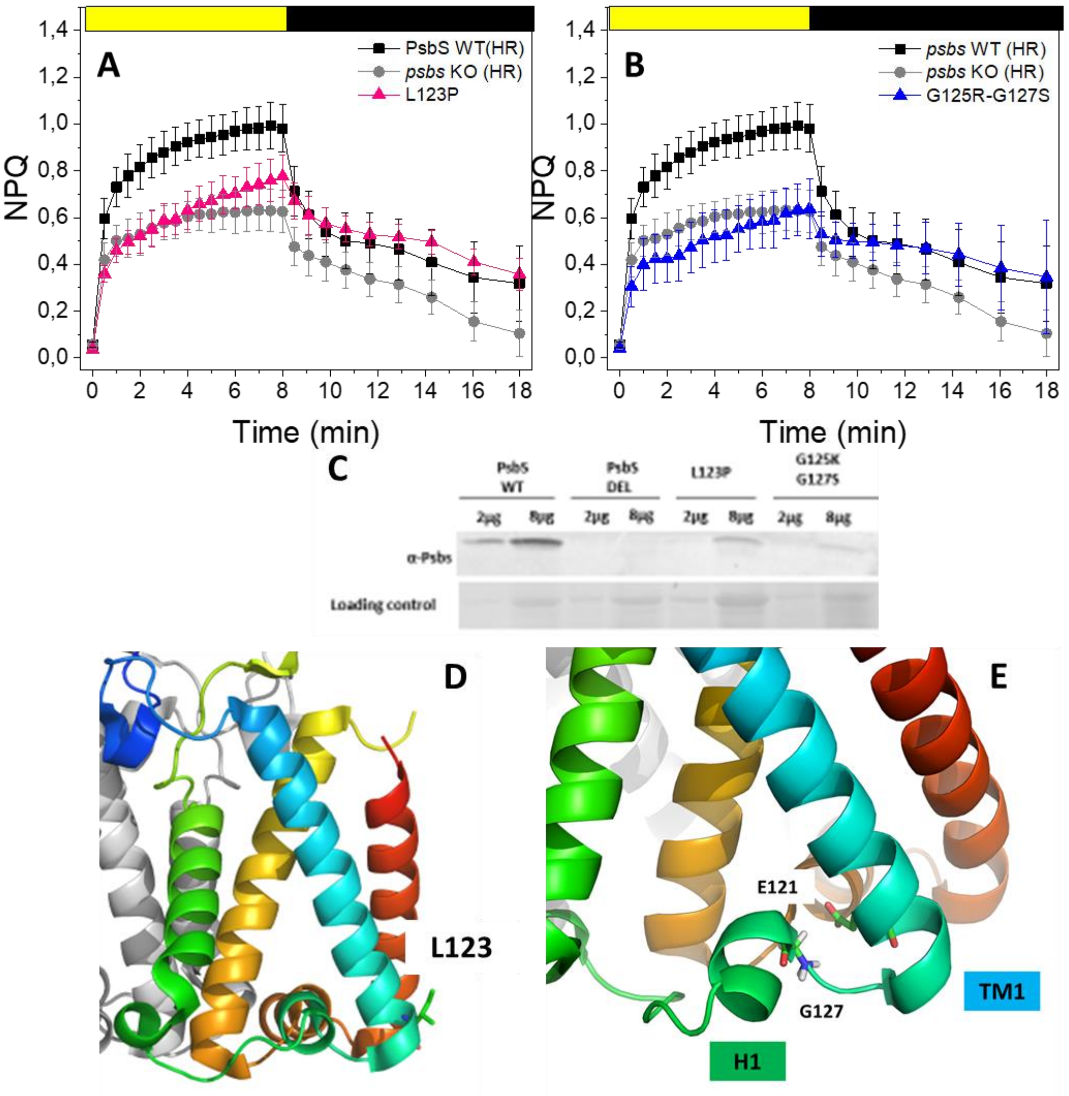


**Figure S10. Mutants with altered PsbS stability.** A-B) NPQ kinetics of L123P (line #140, pink triangles, A) and G125K-G127S (line#41, blue triangles, B) compared with *psbs* WT (black squares) and *psbs* KO (grey circles). For all plants measurements show average of at least 3 independent biological replicates. C) WB analysis comparing PsbS accumulation in *psbs* WT line, line with deletion in PsbS sequence and L123P, G125K-G127S mutant lines. D) PsbS structure showing L123 position (green) at the end of the first transmembrane helix. E) G127 position in PsbS tri-dimensional structure. G127 is shown in sticks. It is located between the first transmembrane helix and the first amphiphilic loop as indicated in the figure.

**Fig. S11.**

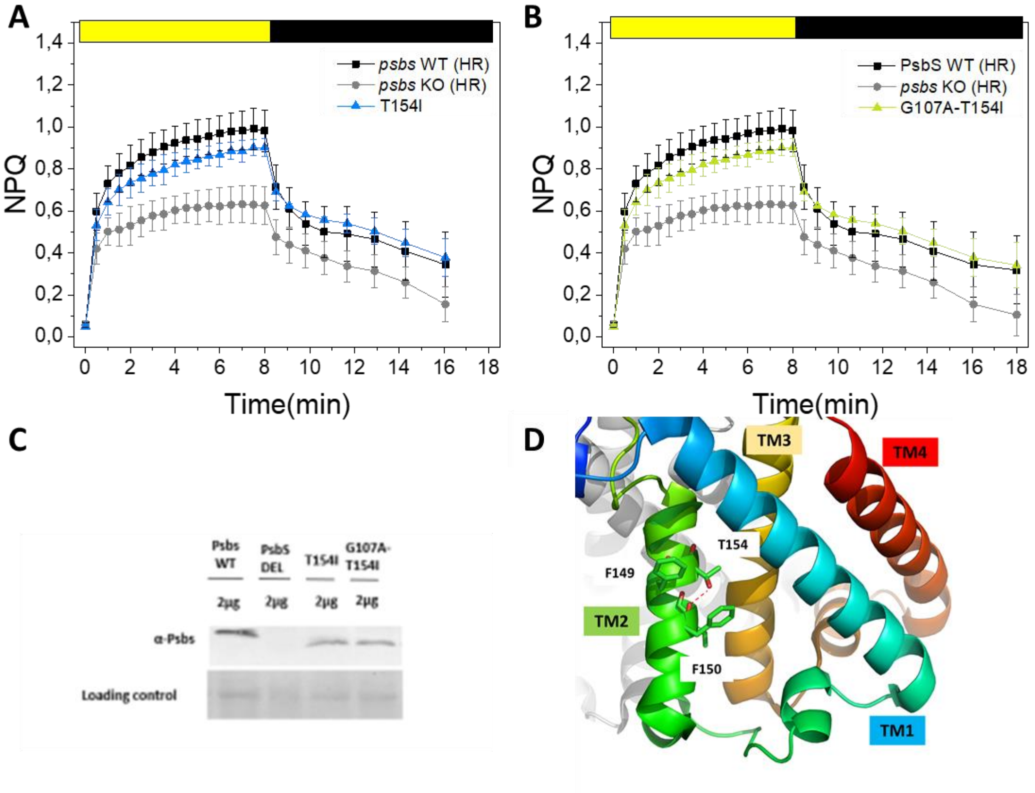


**Figure S11. Mutations causing protein de-stabilization (T154).** A-B) NPQ kinetics of T154I (blue triangles- A) and G107A-T154I (green triangles- B) compared with *psbs* WT (black squares) and *psbs* KO (grey circles). For all plants measurements show average of at least 3 independent biological replicates. C) WB analysis comparing PsbS accumulation in *psbs* WT line, line with deleted PsbS, T154I and G107A-T154I mutant lines. D) Overview of T154 and its interaction with F150. Residues are shown as sticks, polar interactions with the dashed line. Nitrogen and oxygen atoms are coloured in blue and red respectively.

**Fig. S12.**

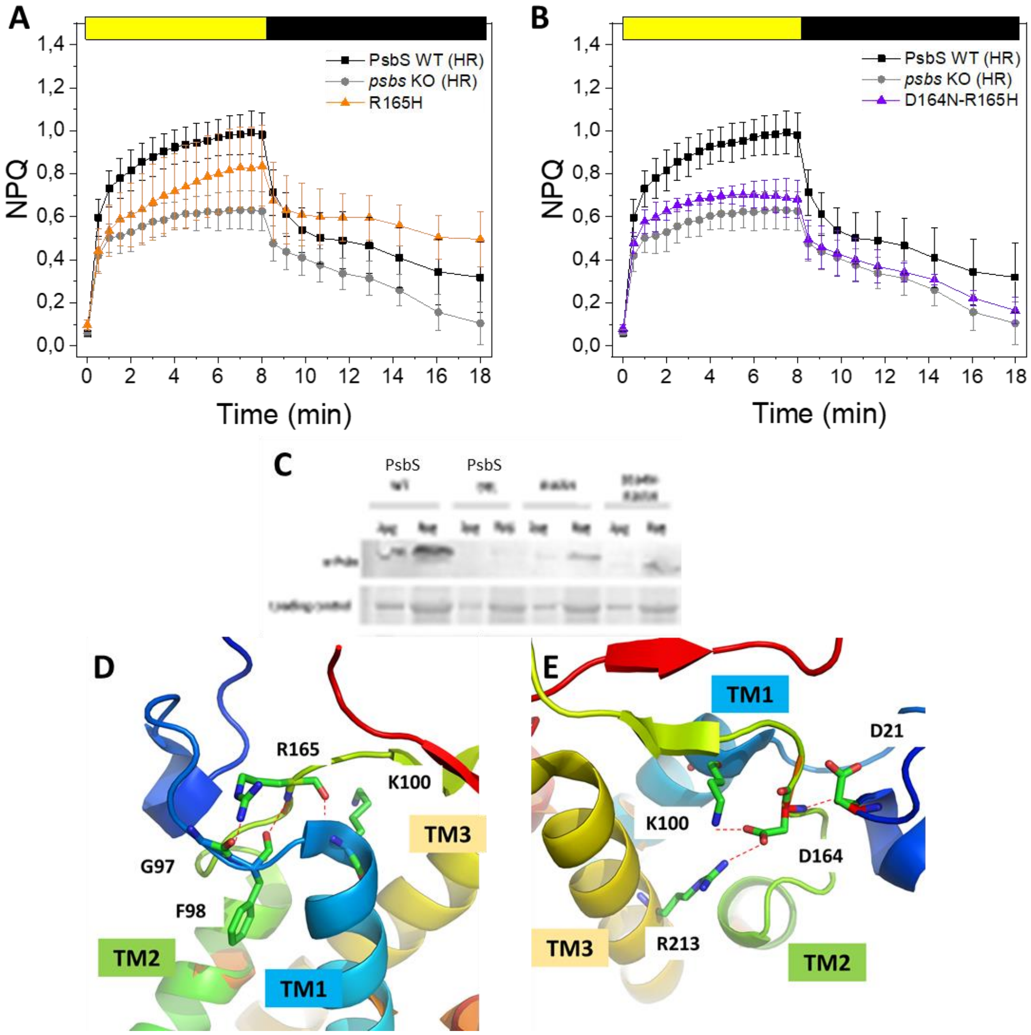


**Figure S12 Mutants with altered PsbS stability.** A) NPQ kinetics of R165H (red triangles, B) and D164N-R165H (orange triangles, B) compared with *psbs* WT (black squares) and *psbs* KO (grey circles). C) WB analysis comparing PSBS accumulation in PsbS WT line, line with deleted PsbS and R165H, D164N-R165H. D-E) General overview of R165 and D164 PSBS tridimensional structure. R165 forms three main interactions; the NH and CO group of the carbon backbone interact respectively with F9898 and K100, while the side chain stabilizes polar interactions with G97. D164 stabilizes polar interactions with K100 and R213 while the NH in the backbone is interacting with D21. Residues are shown as sticks, nitrogen and oxygen atoms are coloured in blue and red respectively. Polar interactions are represented with a red dashed line. Different TMs are shown in different colours.

**Fig. S13.**

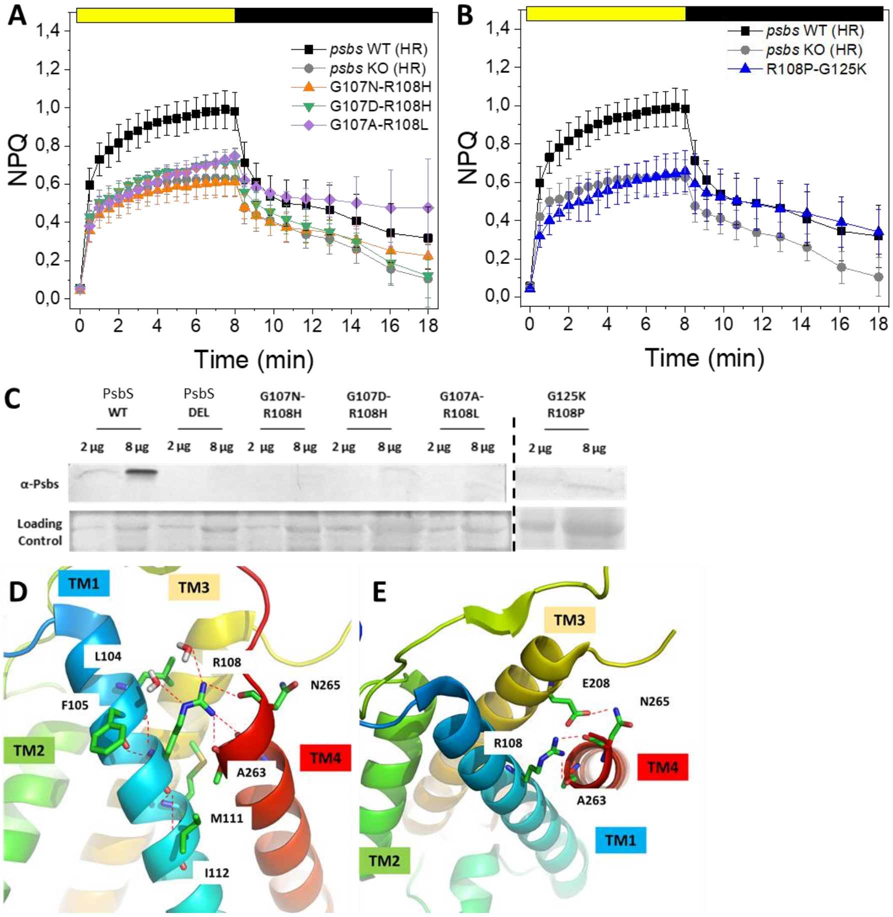


**Figure S13. Mutations causing protein de-stabilization (R108).** A) NPQ kinetics of G107N-R108H (line #197, orange triangles), G107D-R108H (line #198, green triangles), G107A-R108L (line #199, violet triangles). B) NPQ kinetics of G125K-R108P (line #126, blue triangles) compared with *psbs* WT (black squares) and *psbs* KO (grey circles). For all plants measurements show average of at least 3 independent biological replicates. C) WB analysis comparing PSBS accumulation in PsbS WT line, line with deleted PsbS and G107N-R108H, G107D-R108H, G107A-R108L, R108P-G125K. D-E) Overview of R108 position and polar contacts in tridimensional PSBS structure. R108 is located in TM1. Its backbone interacts with L104, F105, M11 and I112. its side chain instead interacts with two water molecules as well as A263 and N265 from TM4. Residues are shown as sticks, polar interactions with the dashed line. Nitrogen and oxygen atoms are coloured in blue and red respectively.

**Fig. S14.**

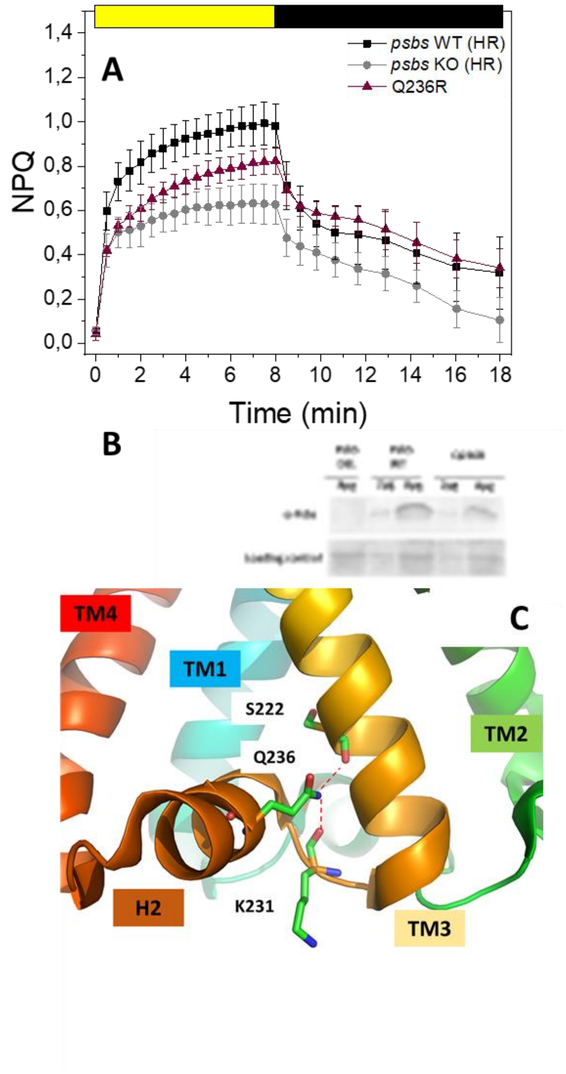


**Figure S14. Mutations causing protein de-stabilization (Q236).** A) NPQ kinetics of Q236R (line#196#205), brown triangles) compared with *psbs* WT (black squares) and *psbs* KO (grey circles). For all plants measurements show average of at least 3 independent biological replicates. B) WB analysis comparing PsbS accumulation in PsbS WT line, line with deleted PsbS and Q236R. C) Overview of Q236 position and polar contacts in tridimensional PsbS structure Q236 is located in the amphiphilic helix H2 in the luminal side. It interacts with K231 and S222 backbones. S222 is not conserved but its interaction involves the backbone NH and thus should not be altered by the residue difference. Residues are shown as sticks, polar interactions with the dashed line. Nitrogen and oxygen atoms are coloured in blue and red respectively.

**Table S1.**

| Name | 5'-3' Sequence |
|---|---|
| sgRNA#5 | ACGGCCAACGAAGAGCTCGTTGG |
| sgRNA#57 | ACGGTCACCCAAAGCACCGATGG |
| sgRNA#176 | TTCACCCTGTTGGGAGCCATCGG |
| sgRNA#180 | GGCGAGATCATTACAGGGAAGGG |
| sgRNA#396 | GGCGAGATCATTACAGGGAAGGG |
| sgRNA#426 | GCCCAACTGAACATTGAAACGGG |
| sgRNA#507 | GCGGCTGTAAACCCCGGAACTGG |

| Name | 5'-3' Sequence |
|---|---|
| PpPsBS-F1 | ATCAAAATACATCATCGGAGA |
| PpPsBS-R1 | AAATAGTAGGGTAGCCAACAT |
| PpPsBs-F2 | CTTCAACCTTGGCCTTGCCCT |
| PpPsBs-R2 | CTTCAACCTTGGCCTTGCCCT |

**Table S1 : Sequences of sgRNAS and primers used in this study used in this study**

**Table S2.**

| | | TM1 | | TM2 | | TM3 | | TM4 |
|---|---|---|---|---|---|---|---|---|
| **line #** | **protein variant** | **sgRNA# 5** | **sgRNA# 57** | **sgRN A#180** | **sgRNA# 176** | **sgRNA #396** | **sgRNA# 426** | **sgRNA# 507** |
| **102** | WT | | | | | silent | | |
| **10** | G125K | | G125K | | | | | |
| **14** | G125E | | G125E | | | | | |
| **23** | G125K | | G125K | | | | | |
| **30** | G125K | | G125K | | | | | |
| **40** | G125K | | G125K | | | | | |
| **51** | G125K | | G125K | silent | | silent | | |
| **57** | G125K | | G125K | | | silent | | |
| **74** | G125K | | G125K | | | | | |
| **85** | G125E | | G125E | | | | | |
| **86** | G125K | | G125K | | | | | |
| **89** | G125K | | G125K | | | | | |
| **111** | G125K | | G125K | silent | | silent | | |
| **112** | G125K | | G125K | silent | | | | |
| **113** | G125K | | G125K | | | | | |
| **129** | G125A | | G125A | | | | | |
| **147** | G125K | | G125K | | | | | |
| **151** | G125K | | G125K | | | | | |
| **152** | G125K | | G125K | | | | | |
| **175** | G125K | | G125K | | | | | |
| **140[a]** | L123P | | L123P | | | | | |
| **193** | F153L | | | F153L | | | | |
| **206** | T154I | | | T154I | | | | |
| **194** | R165H | | | | R165H | | | |
| **195[a]** | I227V | | | | | I227V | | |
| **203[a]** | I227V | | | | | I227V | | |
| **204[a]** | I227V | | | | | I227V | | |
| **196[a]** | Q236R | | | | | | Q236R | |
| **205[a]** | Q236R | | | | | | Q236R | |
| **17** | G95S-G107D | G95S-G107D | | | | | | |
| **41** | G125R-G127S | | G125R-G127S | silent | | | | |
| **49** | G125K-F153L | | G125K | F153L | | silent | silent | |
| **53** | G107A-T154I | G107A | | T154I | | silent | silent | silent |
| **104** | G125K-F153L | | G125K | F153L | | silent | silent | |
| **126** | R108P-G125K | R108P | G125K | | | | | |
| **133** | G125L-F153L | | G125L | F153L | | | | |
| **197** | G107N-R108H | G107N-R108H | | | | | | |

| | | | | | | | | |
|---|---|---|---|---|---|---|---|---|
| **198** | G107D-R108H | G107D-R108H | | | | | | |
| **199** | G107A-R108L | G107A-R108L | | | | | | |
| **200** | D164N-R165H | | | | D164N-R165H | | | |
| **19** | G125K-L155V-G225R | | G125K | L155V | | G225R | silent | |
| **73** | G107D-R108H-Q236E | G107D-R108H | | | | silent | Q236E | |
| **153** | G125K-T154I-Q236E | | G125K | T154I | | | Q236E | |
| **186** | G107D-R108H-G125K | G107D-R108H | G125K | | | | | |
| **121**[b] | G107N-R108H-G125K-Q236R | G107N-R108H | G125K | silent | | | Q236R | |
| **167** | G107A-R108L-G125E-F153L-R165H | G107A-R108L | G125E | F153L | R165H | silent | | |
| **188** | G107A-R108H-G125K-A262G-A263V | G107A-R108H | G125K | silent | | silent | | A262G-A263V |
| **63** | Q236* | | G125K | | | silent | Q236* | |
| **69** | L237* | | G125R | F153L | | silent | L237* | |
| **80** | L237* | G107N-R108H | G125K | F153L | | silent | L237* | |
| **87** | Q236* | G107N-R108H | G125K | | | silent | Q236* | |
| **100** | L237* | G107D-R108H | G125K | | R165P | silent | L237* | |
| **107** | A235V-Q236* | | G125K | | | silent | A235V-Q236* | |
| **122** | G125* | | G125* | | | | | |
| **128** | Q236* | | G125E | silent | | | Q236* | |
| **131** | E135* | R108D | E135* | | | | | |
| **135** | Q236* | G107A-R108L | G125E | silent | | silent | Q236* | |
| **163** | Q236* | | G125A | | | silent | Q236* | |
| **166** | A235V-Q236* | | | | | silent | A235V-Q236* | |
| **178** | G125* | | G125* | | | | | |
| **207** | Q236* | | | | | | Q236* | |
| **208** | Q236* | | | | | | Q236* | |
| **209** | Q236* | | | | | | Q236* | |
| **210** | Q236* | | | | | | Q236* | |
| **211** | L237* | | | | | | L237* | |
| **201** | ΔL147-L155 (9 aa deletion) | | | ΔL147-L155 | | | | |
| **202** | ΔG225-T241 (17 aa deletion) | | | | | ΔG225-T241 | | |
| **142** | ΔG107-V264 (158 aa deletion) | ΔG107-V264 | | | | | | |

**Table S2. List of PsbS protein variants identified in edited plants.** Lines #10-188 were obtained using Multiplex strategy and Lines #193-211 using Simplex strategy. Most lines were edited by CBE but in the lines marked with [a] Lines were edited by ABE or [b] Lines were edited by CBE and ABE. The color gradient from light to dark blue corresponds to 1 to 5 AA changes in the PsbS protein. Lines with premature stop codon or in frame deletions are in orange as lines with small deletions identified from sequencing. Lines used for fine phenotyping analysis are in bold (35 edited lines corresponding to 26 different PsbS variants; 3 lines with in frame deletions, 4 lines deleted non in frame, 6 lines with WT PsbS sequence). Type or paste caption here. Create a page break and paste in the Table above the caption.

**Table S3.**

**List of residues changed in PsbS protein variants and their corresponding aminoacid in spinach (also mature protein) and Arabidopsis.**

.

| **Edited residues** | | | |
|---|---|---|---|
| ***P.patens*** | ***S.oleracea*** | ***S.oleracea* mature protein (Structure)** | ***A.thaliana*** |
| G107 | G103 | G41 | G94 |
| R108 | R104 | R42 | G95 |
| L123 | I119 | I57 | L110 |
| G125 | G121 | I59 | G112 |
| G127 | G123 | I61 | G114 |
| T128 | I124 | I62 | I115 |
| E135 | E131 | E69 | E122 |
| F153 | F149 | F87 | F140 |
| T154 | T150 | T88 | T141 |
| L155V | L151 | L89 | L142 |
| D164 | D160 | D98 | D151 |
| R165 | R161 | R99 | R152 |
| G225 | G220 | I158 | G211 |
| I227 | I222 | I160 | I213 |
| A235 | A230 | A165 | A201 |
| Q236 | Q231 | Q169 | Q202 |
| L237 | L232 | L170 | L203 |
| A262 | A257 | A195 | A248 |
| A263 | A258 | A196 | A249 |
